# Rapid and prolonged response of oligodendrocyte lineage cells in standard acute cuprizone demyelination model revealed by *in situ* hybridization

**DOI:** 10.1101/2023.11.22.568377

**Authors:** Yuehua He, Hua Xie, Liuning Zhang, Yuanyu Feng, Yu Long, ZhengTao Xu, Yanping Zou, Wei Zheng, Shuming Wang, Yongxiang He, Jiong Li, Lin Xiao

**Affiliations:** Key Laboratory of Brain, Cognition and Education Sciences of Ministry of Education; Institute for Brain Research and Rehabilitation, Guangdong Key Laboratory of Mental Health and Cognitive Science, and Center for Studies of Psychological Application, South China Normal University, Guangzhou 510631, China

**Keywords:** Cuprizone, demyelination, remyelination, Multiple sclerosis, oligodendrocyte

## Abstract

Dietary administration of a copper chelator, cuprizone (CPZ), has long been reported to induce intense and reproducible demyelination of several brain structures such as the corpus callosum (CC) in mice, followed by spontaneous remyelination after drug withdrawal. Despite the widespread use of CPZ as an animal model for demyelinating diseases such as multiple sclerosis (MS), the mechanism by which it induces demyelination and then allows robust remyelination is still unclear. An intensive mapping of the oligodendrocyte (OL) lineage cell dynamics during the de-and remyelination course would be of particular importance for a deeper understanding of this model. Here, using a panel of OL lineage cell markers as *in situ* hybridization (ISH) probes, including *Pdgfra, Plp, Mbp, Mog, Enpp6*, combined with immunofluorescence staining of CC1, SOX10, we provide a detailed dynamic profile of OL lineage cells during the entire course of the model from 3.5 days, 1, 2, 3, 4,5 weeks of CPZ treatment, i.e. the demyelination period, as well as after 1, 2, 3, 4 weeks of recovery (drug withdrawal) from 5 weeks of CPZ treatment, i.e. the remyelination period. The result showed an unexpected early death of mature OLs and response of OL progenitor cells (OPCs) in vivo upon CPZ challenge, and a prolonged upregulation of myelin-forming OLs compared to the intact control even 4 weeks after CPZ withdrawal. These data may point to the need to optimize the timing windows for the introduction of pro-remyelination therapies in demyelinating diseases such as MS, and may serve as a basic reference system for future studies of the effects of any intervention on demyelination and remyelination using the CPZ model.

## Introduction

Myelin in the central nervous system (CNS) is produced by oligodendrocytes (OLs). Myelin sheaths are specialized membrane extensions of OLs that tightly wrap axons with a multi-layered, concentric, ring-like structure. It insulates the axon and allows for rapid saltatory conduction of action potentials, greatly increasing the efficiency of the nervous system while being highly energy efficient (Li and Richardson, 2016). The myelination of axons was a truly momentous event in the history of biological evolution. The complete myelin structure evolved and first appeared in cartilaginous fish ∼600 million years ago, and was maintained from then on in all jawed vertebrates (Li and Richardson, 2016; Nave and Trapp, 2008). Demyelinating diseases of the CNS are a group of neural disorders characterized by the progressive and eventually irreversible loss of OLs and myelin structures, which can lead to a range of neurological and psychiatric dysfunctions. While recent years has witnessed a growing body of evidences showing demyelinating damages as a critical brain pathological change in neurodegenerative diseases such as Alzheimer’s disease (Wang et al., 2018) and various neuropsychiatric disorders such as major depression (Williams et al., 2019), schizophrenia (Miyata et al., 2015), and autism (Shen et al., 2018), the most representative and widely studied demyelinating disease is multiple sclerosis (MS).

First clinically described over 120 years ago, MS is now generally accepted be an inflammatory, autoimmune demyelinating disease of the CNS in which an adaptive immune response targets unknown CNS antigens, resulting in myelin damage, and subsequent impaired conduction of action potential and axonal degeneration (Friese et al., 2014; Frohman et al., 2006; Noseworthy et al., 2000). Patients with MS can present with a variety of neurological dysfunctions, including ataxia, dexterity loss, myoclonus, spasticity, paraparesis or hemiparesis, vision loss and cognitive deficits, depending on the different pathways involved. Despite more than a century of research, the precise etiology and underlying pathophysiological mechanisms of MS remain poorly understood and clinical strategies are limited (Dobson and Giovannoni, 2019; Walton et al., 2020). Because of the different cellular mechanisms that characterize the different stages of MS, it is not possible to model the disease as a whole with a single model. To date, four types of animal models have been developed to model the different aspects of MS, including: genetic myelin mutations, virus-induced demyelination, experimental autoimmune encephalomyelitis (EAE), and toxin provoked demyelination. Among these, the most widely used models are EAE and cuprizone (CPZ) toxicity (Hooijmans et al., 2019; Vega-Riquer et al., 2019; Zirngibl et al., 2022). Although EAE is thought to mimic the lesion formation and inflammatory injury features of relapsing-remitting MS, the most common clinical representation of MS. The widespread immune response and the random appearance of demyelinating lesions make it difficult to disentangle the intrinsic immune relevance of the CNS to disease progression and remyelination (Lassmann and Bradl, 2017; Ransohoff, 2012). Therefore, toxin-based animal models such as the CPZ-induced demyelination, which limit random lesion formation and peripheral immune cell infiltration, have been employed by researchers as an alternative choice (Sriram and Steiner, 2005).

Despite the widespread use of CPZ, the mechanism by which it induces demyelination and allows spontaneous remyelination without damaging to other types of neural cells in the brain, is still unclear. Copper, one of the most essential trace elements for the human body, is involved in a number of important biological processes, such as serving as a component of metalloenzymes and acting as an electron donor or acceptor, which is critical for cellular energy production (Stern et al., 2007). Early evidence suggests that CPZ-induced mitochondrial dysfunction and the subsequent energy and metabolic disturbances may be the primary cause of OL death and myelin loss. As many of these studies have used different strains, sexes and ages of mice, it has been difficult to interpret results between experiments, highlighting the need for a more standardized procedure (Zirngibl et al., 2022). The recent introduction of 0.2% (w/w) protocol in young adult C57BL/6 male mice as a standardized model has significantly improved the situation and led to a deeper understanding of the mechanism for CPZ-induced demyelination, e.g. the role of astrogliosis and microgliosis and the various local inflammatory factors released by them (Vega-Riquer et al., 2019; Zirngibl et al., 2022). However, a whole process mapping of the dynamics of OL lineage cells during both the demyelination and remyelination phases of the standardized CPZ model is still lacking (Jhelum et al., 2020).

In this study, taking advantage of a panel of *in situ* hybridization (ISH) probes targeting the marker genes for different stages of OL lineage cells, including *Pdgfra, Plp, Mbp, Mog,* and especially the recently reported maker gene for newly forming OLs, namely *Enpp6*, with the combination of immunofluorescence staining of CC1, SOX10, we provide a temporally detailed dynamic profile of OL lineage cells in the currently most commonly used standardized acute CPZ demyelination model of 5 weeks feeding followed by 4 weeks recovery. Our results show a rather rapid and prolonged response of OL lineage cells to the CPZ challenge in this model.

## Materials and Methods

### Animals

Male C57BL/6 mice (6 weeks old) were acquired from Guangzhou Yancheng Biotechnology Co, LTD and housed with a 12 h dark/12 h light cycle, at a constant temperature of 24 ± 1 °C and a relative humidity of 55±15%. Food and water were available ad libitum. All mice were acclimated to laboratory conditions for 1 week before starting the experiments, followed by 1 week of acclimatization to normal powder chow before the starting of experiments at 8 weeks of age. All animal experiments were approved by the Animal Care and Use Committee of South China Normal University and followed the guidelines published in the National Institutes of Health Guide for Care and Use of Laboratory Animals.

### Cuprizone treatment and tissue processing

CPZ treatment of animal were carried out as previously described with some modifications (Li et al., 2013; Xiao et al., 2012) Briefly, mice were fed with on rodent chow containing 0.2% (w/w) cuprizone (bis-cyclohexanone oxaldihydrazone, Sigma #14690-100G) for different durations up to 5 weeks before sampling for the detection of demyelination and tracing of the OL lineage cell dynamics in the demyelination courses. For analysis of the remyelination courses, mice were fed with the same dose of CPZ for 5 weeks and then returned to normal chow without CPZ for recovery from demyelination for 1-4 weeks before sampling. For brain tissue sampling, animals were sacrificed at different time points (3.5d, weeks 1, 2, 3, 4, and 5 for demyelination; weeks 6, 7, 8 and 9 for remyelination). Under deep anesthesia with pentobarbital sodium (50 mg/kg, i.p.) C57BL/6 mice in each group were perfused intracardially with 0.01M RNase-free phosphate buffered saline (PBS, pH 7.4), followed by 4% (w/v) paraformaldehyde (PFA; Sigma). Brains were removed, post-fixed in the same fixative overnight at 4°C, and cryoprotected in 20% (w/v) sucrose (Sigma) overnight followed by 30% (w/v) sucrose overnight, embedded in OCT medium and stored at -80°C until use. Under RNase-free conditions, serial coronal sections of 25 μm thickness were cut with a cryostat and collected in 6-well plates containing 0.01 M RNase-free PBS. All solutions were prepared with DEPC treated water. The middle region of the corpus callosum in coronal sections approximating to bregma +0.74 mm to -0.82 mm according to Paxinos and Franklin’s atlas were used for all experimental analysis.

### Western blot

Animals were given intraperitoneal injection of 50 mg/kg sodium pentobarbital, and brains were rapidly removed and frozen. Corpus callosum tissue was isolated from the brains of differently treated mice and sampled by homogenization in RIPA lysis buffer (Beyotime Biotechnology) supplemented with a protease inhibitor (Sigma). Total protein concentration was measured using a BCA assay kit (TIANGEN). Then 10 mg of protein per sample was separated on 12% SDS-PAGE gels (Sangon Biotech) and subsequently transferred to PVDF membrane (Millipore, Temecula) using sodium dodecyl sulfate-polyacrylamide gel electrophoresis and Western blotting techniques. Membranes were blocked with 5% milk in 0.1M TBST and probed with primary antibody: GAPDH (Mouse, Abcam G8795,1:1000) was used as an internal control for the concentration of protein loaded, while anti-MBP (Rat, BioRad MCA4095,1:1000) was used to detect MBP. Blots were then washed and incubated with HRP-conjugated IgG (Mouse, Abcam ab6789,1:5000) and (Rat, Bioss DS-0293G,1:5000), the immunoreactive band was visualized using an ECL detection kit (Beyotime Biotechnology). Scanning densities were normalized to control and expressed as relative folds of control. Results were from three independent experiments.

### LFB-PAS Staining and quantification

Luxol fast blue (LFB) staining of the myelin was performed as previously described (Xiao et al., 2012). Briefly, sections were defatted in 1:1 ethanol/chloroform solution for 2h and then dehydrated through 95% ethanol (Sangon Biotech) washes. Then incubated in a 0.1% LFB solution (Sigma #1002798699) more than 8 hours at 56°C. Sections were rinsed two to three times in distilled water to remove excess stain, dehydrated through graded ethanol washes and differentiated in 0.05% lithium carbonate (Sigma #102076768) until grey and white matter could be distinguished macroscopically. Sections were oxidized in 0.5% periodic acid (PA) solution (Sigma #102025710) for 5 min, rinsed in distilled water, and stained in Schiff reagent (Sigma #1.09033.0500) for 15 min then washed in lukewarm water for 5 min. The nuclei were stained with hematoxylin (Sigma #E607317) for 30 s, followed by washing in distilled water. Subsequently, xylene was used to make the slices transparent and neutral gum to seal them (Carriel et al., 2017). Sections can be stained blue for myelin by using LFB, while periodic acid-Schiff’s (PAS) stains microglia/macrophages and demyelinated axons pink. To evaluate the extent of demyelination and remyelination, all sections were scored by two blinded observers on a scale from 0 (complete myelination) to 3 (complete loss of myelin), with a score of 1 or 2 corresponded to one-third or two-thirds myelination of CC fibers. The intermediate score reflects the intermediate proportion of demyelinated fibers. Representative section for each score is shown in supplemental figure 1.

### Immunohistochemistry, microscopy imaging and cell counts

Coronal cryosections (25μm) of the brain were collected and processed as floating sections. Sections were then washed three times in PBS, treated with citrate for 20 min to repair the antigens, and washed again three times in PBS. Then blocked with 10% goat serum in PBS with 0.3% TritonX-100 for 2 hours, and immediately incubated with a primary antibody in a blocking solution at 4°C overnight. Primary and secondary antibodies were diluted in blocking solution (0.3% (v/v) Triton X-100, 10% (v/v) FBS in PBS). Sections were then washed three times in PBS and incubated with secondary antibodies (Alexa Fluor 488/555, Abcam, 1:500) and DAPI for 2 hours at room temperature. After three rinses in PBS, the sections were mounted on glass slides. Primary antibodies were anti-SOX10 (guinea pig, Hangzhou Normal University,1:200), monoclonal CC1 (mouse, Calbiochem OP80, 1:200). Low-magnification (20x objective) confocal images were collected using a Zeiss laser Airyscan confocal microscope (LSM 900) as Z stacks with 1-μm spacing, using standard excitation and emission filters for DAPI, FITC (Alexa Fluor 488), TRITC (Alexa Fluor 555) excitation and emission filters were used. Cells were counted blindly by two independent observers in non-overlapping fields of coronal sections of the corpus callosum between the dorsolateral corners of the lateral ventricles (six fields per section, three sections from each of three or more mice of a given experimental group).

### In situ hybridization

*In situ* hybridization (ISH) was performed on fresh frozen coronal brain sections (25 µm) according to protocols that are available at http://www.ucl.ac.uk/∼ucbzwdr/Richardson.htm. Briefly, digoxigenin (DIG)-labeled RNA probes were transcribed in vitro from cloned cDNAs for mouse *Pdgfra, Enpp6, Plp, Mbp* and *Mog* as previously described (Huang et al., 2018; Jolly et al., 2016; Xiao et al., 2016). Brain slices were incubated with DIG-labeled RNA antisense probes overnight at 64∼66°C, depending on the probes. Slices were then washed 3 times with MABT buffer, followed incubation with blocking solution for 2 hours before incubation with alkaline phosphatase (AP)-conjugated anti-DIG Fab fragments antibody (1:1000, Roche, #11093274910) overnight at 4°C. The targeted mRNA signals were visualized using NBT (nitroblue tetrazolium, Roche, #11383213001)/BCIP (5-bromo-4-chloro-3-indolyl phosphate, Roche, #11383221001) color development substrate via AP activity. Optic images were taken by the LEICA Camera microscope (MD4B). Cell counts were carried out in the same way as above described.

### Quantification and statistical analysis

All data presented are expressed as arithmetic mean± SEM. Cell counts were performed in as blinded fashion. No statistical methods were used to predetermine sample sizes, but our sample sizes are similar to those typically reported in the field. Differences between groups were analyzed using two-tailed unpaired Student’s t tests with GraphPad Prism 8.0 software. The statistical significance of the difference of each group compared to the control is indicated (\**p*≤0.05; \*\**p*≤0.01; \*\*\**p*≤0.001; ns, no significant difference). Exact statistical p-values are given in the results text.

## Results

### Confirmation of de- and subsequent remyelination in the standardized CPZ model in C57BL/6 mice

To standardized the CPZ model, we used young adult male C57BL/6 mice aged 8-weeks for CPZ feeding at a dose of 0.2%(w/w) in all experiments (Hiremath et al., 1998).Before tracking OL lineage cell dynamics in this standardized CPZ model, we first confirmed and mapped out a detailed time course with mainly 1-week intervals for the CPZ-induced demyelination and subsequent remyelination after drug withdrawal, with the whole course lasting up to 9 weeks, as shown in the diagram in (Fig.1A). We used the well-documented LFB-PAS staining and scoring protocol (Hiremath et al., 1998) to show the myelination degree of mice at different time points under their respective treatments. We analyzed the middle part of the CC as shown in (Fig.1B), which is the most commonly referred and well-described brain region reproducibly affected in the CPZ model (Hiremath et al., 1998; Vega-Riquer et al., 2019). Consistent with previous reports, our results showed that there was a slight decrease in myelination score (demyelination) at 2 weeks after CPZ treatment. Demyelination became significant at 3 weeks, and progressed strongly during the 4^th^ and 5^th^ weeks of CPZ feeding, reaching the bottom (i.e. complete demyelination) at 5 weeks. Remyelination occurred rapidly and robustly as soon as 1-week after CPZ withdrawal, and the myelination scores reached levels similar to intact controls after 3-4 weeks of recovery (Fig.1G-H). For detailed myelination scores at each time points: 3.5 days (CPZ, 3 vs. Control, 3); 1w ( CPZ, 2.786±0.178 vs. Control, 3; p=0.0833); 2w (CPZ, 2.75±0.13 vs. Control, 3; p=0.0152); 3w (CPZ, 2.25±0.30 vs. Control, 3; p=0.0079); 4w (CPZ, 0.52±0.22 vs. Control, 3; p<0.0001); 5w (CPZ, 0.36±0.34 vs. Control, 3; p<0.0001); 5+1w (CPZ, 1.52± 0.40 vs. Control, 3; p=0.0007); 5+2w (CPZ, 1.81±0.08 vs. Control 3; p<0.0001); 5+3w (CPZ 2.58±0.45 vs. Control, 3; p=0.1589); 5+4w (CPZ, 2.63±0.38 vs. Control, 3; p=0.134). We also collected a protein lysis sample from the fresh CC tissue for WB analysis at time of peak demyelination, which showed a dramatic decrease in the myelin basic protein (MBP) levels in 5-week CPZ-treated mice (CPZ, 0.60±0.16 vs. Control, 1±0.03; p=0.0227) (Fig.1C-D). In addition, immunofluorescence staining also showed a dramatic decrease in MBP signals after 5 weeks CPZ treatment (CPZ, 23.47±2.67 vs. Control, 95.47±2.25; p<0.0001) (Fig.1 E-F). These results confirmed that in our standardized model, 5 weeks of 0.2% CPZ feeding in young adult C57BL/6 mice induced profound acute demyelination in the CC region followed by almost complete remyelination after 4 weeks of recovery by CPZ withdrawal. We then set out to map the dynamics of OL lineage cells in this standardized model.

**Fig. 1.**
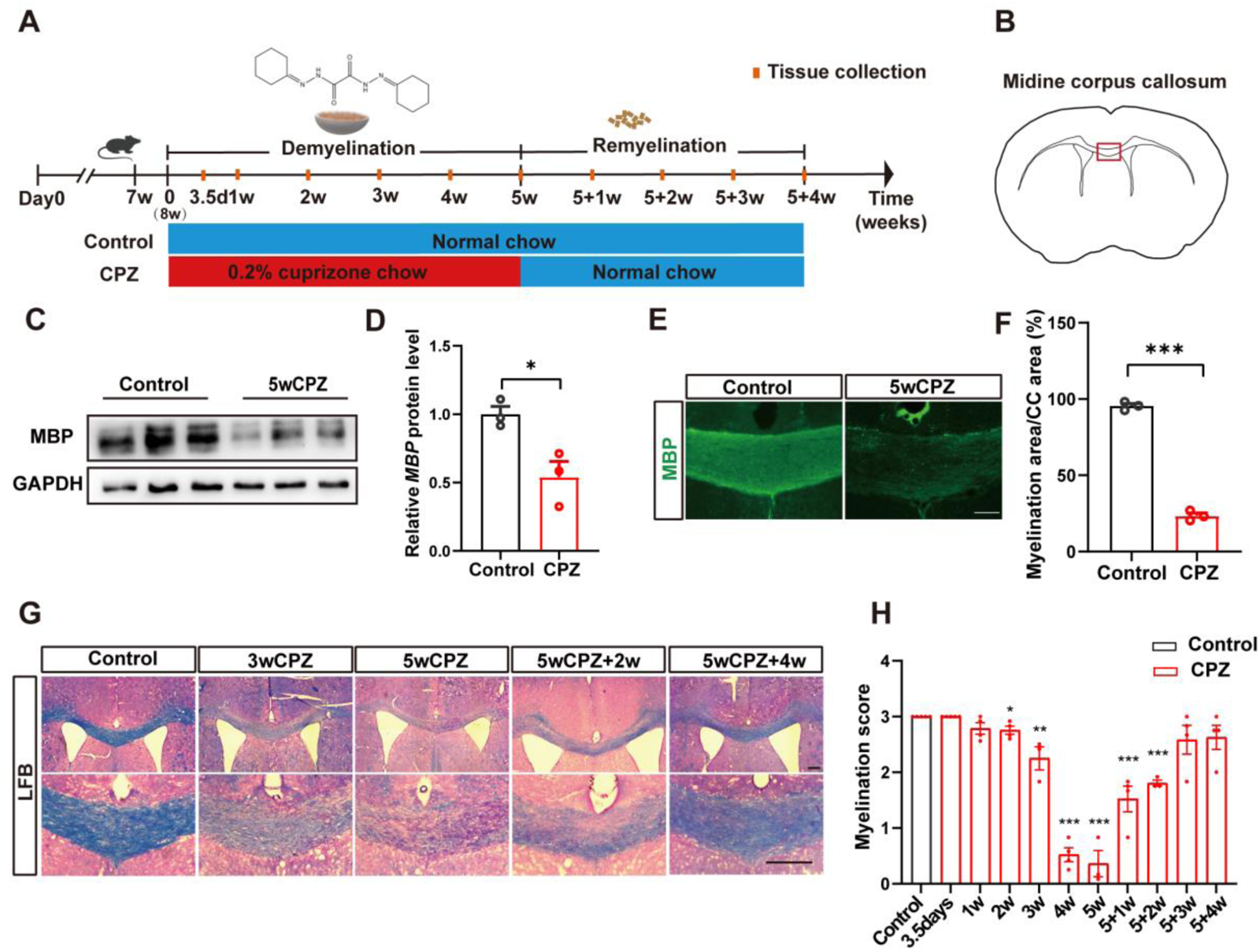
Establishment of standard CPZ de- and remyelination model. (A) Representation of the cuprizone exposure protocol. Normal chow (blue); 0.2% CPZ treatment (red); time point of tissue collection (orange line); (B) The midline corpus callosum was evaluated. (C) Representative Western blotting for corpus callosum of mice fed with 5w or without CPZ. (D) Quantification of MBP protein level. (n=3/group) (E) Immunofluorescence labeling of MBP. Representative images from control and 5w CPZ-treated mice. (F) Quantification of MBP protein area. (n=3/group) (G) Representative Luxol fast blue staining images from control, CPZ-treated mice 3w, 5w; and 2, 4 weeks after withdrawal of CPZ chow. (H) Quantification of myelin sheath score. (3.5d, n=5/group; 1, 2w, 4w, 5+1w, 5+3w, 5+4w, n=4/group; 3w, 5w, 5+2w, n=3/group). Data are mean± SEM. Asterisks indicate significant difference from mice on the normal chow mice. \**p*<0.05; \*\**p*<0.01; \*\*\**p*<0.001. Two-tailed unpaired t-test. Scale bars: 100 μm; CPZ, cuprizone; MBP, myelin basic protein;

### Dynamics of CC1-positive mature OLs and SOX10-positive OL lineage cells as a whole

Mature myelinating OLs are generated from their precursors, called oligodendrocyte progenitor cells (OPCs), by gradual differentiation that undergo a sequence of intermediate maturation stages with distinct cell markers during CNS development (Bergles and Richardson, 2016; Nishiyama et al., 2009). To track the changes in the number of mature OLs throughout the course of the above standardized CPZ model, immunofluorescence staining was performed using the monoclonal antibody anti-adenomatous polyposis coli (APC) clone CC1, which is the most commonly used tool to specifically label mature OL cell bodies without labeling myelin and is thus very suitable for identifying and counting the OLs in tissue sections (Bin et al., 2016). Sections were simultaneously double stained with antibodies targeting SOX10 (Fig.2A), which is a well-known transcription factor that is specifically expressed by all OL lineage cells of different mature stages in the brain. As shown in Figure 2, the number of CC1 and SOX10 double-positive cells decreased rapidly (as early as 3.5 days) and dramatically (by ∼40% reduction) (CPZ, 2462.63±488.59 cells/mm^2^ vs. Control, 4100.67±410.77cells/mm^2^; p=0.0009) in response to CPZ challenge, and hit bottom at 2 weeks (by ∼93% reduction) (CPZ, 268.61±73.20 cells/mm^2^ vs. Control, 4081.82±391.70 cells/mm^2^; p<0.0001) followed by a rapid recovery to a level near control only 1 week after CPZ withdrawal (5+1w (CPZ, 3253.71±366.94 cells/mm^2^ vs. Control, 3996.42±389.59 cells/mm^2^; p=0.053)), while that of the control remains basically unchanged during the whole 9 weeks’ period (Fig.2B). For detailed cell counts at other time points (in cells/mm^2^): 1w (CPZ, 810.41± 207.82 vs. Control 3759.71±457.94; p<0.0001); 3w (CPZ, 481.95±45.42 vs. Control 3812.25±336.72; p=0.0002); 4w (CPZ, 1035.45±54.81 vs. Control 3738.63±536.23; p=0.0001); 5w(CPZ, 2113.6±235.47 vs. Control, 4111.05±78.70; p=0.0003); 5+2w (CPZ, 4148.72±383.85 vs. Control, 4482.05±256.53; p=0.3649); 5+3w (CPZ, 3480.71±35.84 vs. Control, 3845.74±361.83; p=0.1327); 5+4w (CPZ, 3969.22±341.82 vs. Control, 4268.56±152.06; p=0.2151). By calculating the proportion of CC1^+^/SOX10^+^ mature OLs in the total SOX10^+^ OLs lineage cells pool, it is found that CC1^+^ mature OLs made up >90% of all OLs lineage cells in the control mice at all time points. CPZ treatment rapidly decreased this proportion by ∼12% at 3.5 days (CPZ, 82.31%±0.11%; Control, 93.90%±0.01%; p=0.0682), and drove it to a bottom of ∼47% reduction at 3-week (CPZ, 51.93±0.02% vs. Control, 95.17%±0.01%; p<0.0001). The proportions were then also rapidly recovered to a level near normal control from 5+1w (CPZ, 91.17±0.03% vs. Control, 95.33%±0.01%; p=0.0635) and then on (Fig.2D). The dynamic of the total number of SOX10^+^ OLs lineage cells showed a very similar curve to that of CC1^+^/SOX10^+^ mature OLs (Fig.2C).

**Fig. 2.**
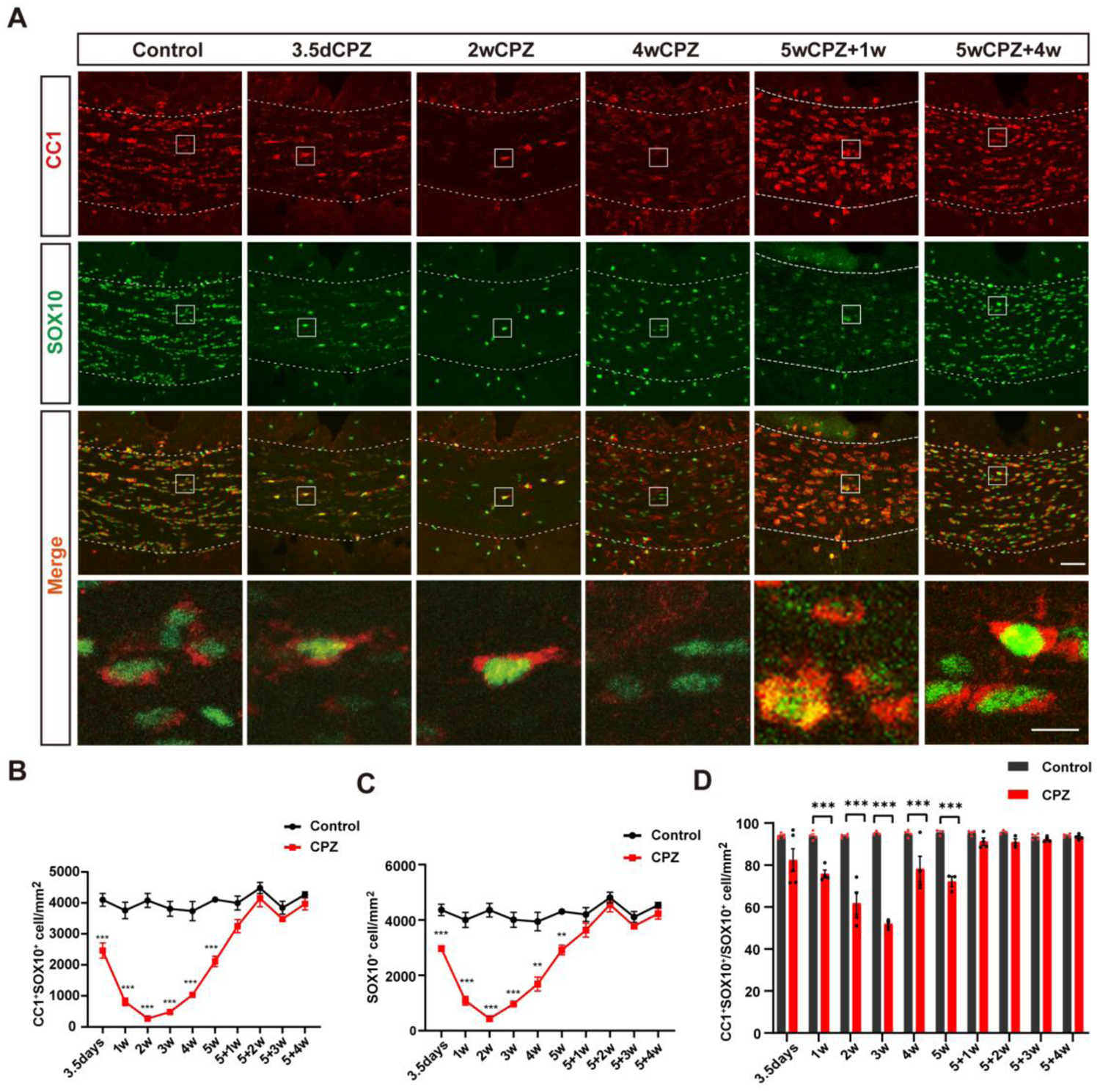
Dynamics of CC1-positive mature OLs and SOX10-positive OL lineage cells as a whole. (A) Double-immunofluorescence labeling of CC1 (red) and sox10 (green) taken from the midline region of the CC at different time points with CPZ chow. Representative images of control, CPZ-treated mice at 3.5d, 2w, 4w; and 1, 4 weeks after withdrawal of CPZ. Images were acquired using confocal laser scanning microscopy(20X) (B)The graph shows plots of the CC1^+^SOX10^+^ cells number in the same region for control (black line) and treatment group (red line) during de-/ remyelination. Note the dynamic changes in the peaks and troughs of oligodendrocyte expression. (C) Graph shows plots of the SOX10^+^ cells number in the same region for control (black line) and treatment group (red line) during de-/ remyelination. (D) Figure shows the proportion of CC1^+^SOX10^+^ cells among SOX10^+^. (3.5d, n=5/group; 1, 2w, 4w, 5+1w, 5+3w, 5+4w, n=4/group; 3w, 5w, 5+2w, n=3/group). Data are mean± SEM. Asterisks indicate significant difference from mice on the normal chow mice. \**p*<0.05; \*\**p*<0.01; \*\*\**p*<0.001. Two-tailed unpaired t test. Scale bar: 50 μm, magnified images scale bar: 10um; CPZ, cuprizone; CC1, mature OL marker; SOX 10, pan-oligodendrocyte lineage cells marker.

### Dynamics of *Pdgfra*-positive OPCs revealed by ISH

Recruitment and proliferation of OPCs often occurred at the demyelinating lesion sites of MS brain (Franklin and Ffrench-Constant, 2008). This process also occurred in several CPZ model protocol settings as previously shown by immunostaining for NG2 or *Pdgfra* (Gudi et al., 2009; Mason et al., 2000), two widely-used OPC cell membrane markers. Platelet-derived growth factor receptor alpha (*Pdgfra*) is a powerful and reliable marker of OPCs in the CNS with high specificity (Pringle NP et al., 1992). To facilitate a clearer and more reliable cell counting to trace the dynamics of OPCs in the current standardized acute CPZ model, we performed an *in situ* hybridization experiments using a previously tested well-functioning probe for *Pdgfra* (Xiao et al., 2016). As shown in Figure 3, similar to the rapid reduction in CC1-positive mature OLs, we also observed a rapid and significant reduction (by ∼52%) (CPZ, 80.51±13.01 cells/mm^2^ vs. Control, 165.36±7.18 cells/mm^2^; p<0.0001) in the number of *Pdgfra*-positive OPCs as early as 3.5 days after CPZ administration, and remained below control up to the 2-week time point (CPZ,108.09±5.37 cells/mm^2^ vs. Control,152.02±18.70 cells/mm^2^, p=0.0079). This decrease was then reversed at week 3 (CPZ,281.59±61.33 cells/mm^2^ vs. Control,155.76±17.71 cells/mm^2^; p=0.0494) and followed by a sharp increase to peak at week 4 at a level of ∼4 folds that of control (CPZ, 628.71±66.91 cells/mm^2^ vs. Control,149.05±17.14 cells/mm^2^; p<0.0001). Following this peak, there was an initial also sharp but then gradual decline towards control, while still remaining ∼67% higher than control 4 weeks after CPZ withdrawal (CPZ, 200.33± 13.65 cells/mm^2^ vs. Control 119.79±14.73 cells/mm^2^; p=0.0004) (Fig.3A, B). The number of *Pdgfra*-positive OPCs in the no-CPZ control mice stayed fairly stable throughout the 9-weeks period. For detailed cell counts at other time points (in cells/mm^2^): 1w (CPZ,102.05± 22.82 vs. Control,172.67±8.56; p=0.0024); 5w (CPZ,336.66±53.55 vs. Control, 124.64±11.53; p=0.0054); 5+1w (CPZ, 261.77±51.83 vs. Control,154.72±19.15; p=0.0153); 5+2w (CPZ, 272.98±29.51 vs. Control 110.77±9.34; p=0.0018); 5+3w (CPZ, 263.97±26.23 vs. Control 150.42±22.01; p=0.0012). This result indicates a significant early response of OPCs to CPZ challenge, with a robust accumulation/proliferation during the course of demyelination and prolonged activity during the remyelination period.

**Fig. 3.**
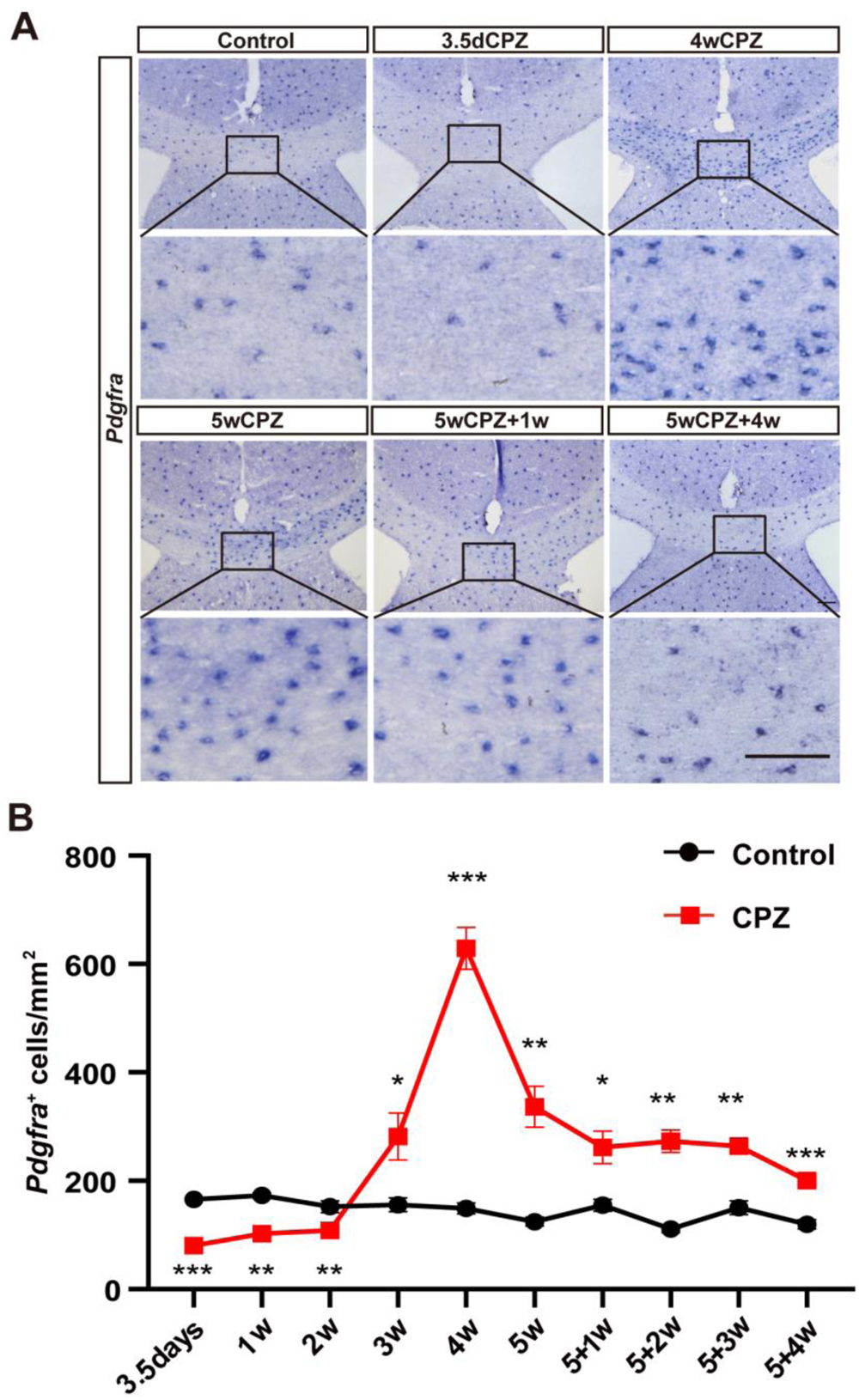
Dynamics of *Pdgfra*-positive OPCs in standard CPZ model. (A) For brain slice samples taken at different weeks, the number of OPCs in the corpus callosum was quantified by *in-situ* hybridization with a *Pdgfra*-specific probe. Representative images from control, CPZ-treated mice 3.5d, 4w, 5w; and 1, 4 weeks after withdrawal of CPZ. (B) Graph showing quantification data of the number of OPCs from control (black line) and experimental (red line) sacrificed animals at different time points. (3.5d, n=5/group; 1, 2w, 4w, 5+1w, 5+3w, 5+4w, n=4/group; 3w, 5w, 5+2w, n=3/group). Data are mean± SEM. Asterisks indicate significant difference from mice on the normal chow mice. \**p*<0.05; \*\**p*<0.01; \*\*\**p*<0.001. Two-tailed unpaired t test. Scale bars: 100um. CPZ, cuprizone; OPCs, oligodendrocyte precursors; *Pdgfra*, platelet-derived growth factor receptor alpha gen, OPCs marker.

### Dynamics of mature OLs revealed by ISH using a panel of myelin-related gene probes

Mature OLs express a number of myelin-related genes that encode proteins critical for the formation of the compacted myelin structure, such as myelin basic protein (MBP), proteolipid protein (PLP) and myelin oligodendrocyte glycoprotein (MOG). These proteins are highly abundant in the white matter of the adult brain and widely distributed in the process/myelin segment of mature OLs while not accumulating in the OL cell body like CC1. To further follow the dynamics of mature OLs in an alternative way to CC1 immunostaining, we performed ISH of the above three myelin related-genes and counted the number of positive cells at different time points throughout the CPZ model. We found a rapid and unexpectedly large decrease in the number of both *Plp-* and *Mog-*positive cells from 3.5 days on after CPZ intake, with only ∼6% (CPZ,180.94±22.85 cells/mm^2^; Control, 2867.97±339.85 cells/mm^2^; p<0.0001) and ∼3% (CPZ, 40.35±19.57 cells/mm^2^ vs. Control,1442.015±128.20 cells/mm^2^; p<0.0001) remained compared to control, respectively (Fig.4, Fig.5). The number of these cells stayed at the lowest level for the following 2 weeks and showed a slight increase from 3 to 5 weeks but still remained below the control. However, this situation was reversed in the remyelination period, there was a sharp and dramatic increase during the first week of recovery after CPZ withdrawal, causing these cell numbers to exceed the control groups, and they maintained at the high level for at least 3 more weeks thereafter (Fig.4B, Fig.5B). For detailed *Plp-*positive cell counts (in cells/mm^2^): 1w (CPZ, 147.64±2.63 vs. Control, 2898.73±203.41; p<0.0001); 2w (CPZ, 144.53±11.45 vs. Control, 2631.40±143.34; p<0.0001); 3w (CPZ, 733.88±115.91 vs. Control, 2519.06± 44.59; p<0.0001); 4w (CPZ, 641.79±76.10 vs. Control, 2614.56±371.26; p=0.0001); 5w (CPZ, 1188.31±314.23 vs. Control, 2475.40±318.67; p=0.0153); 5+1w (CPZ, 2849.87±166.66 vs. Control, 2547.61±90.53; p=0.0328); 5+2w (CPZ, 2317.95± 215.24 vs. Control, 2586.67±322.87; p=0.3829); 5+3w (CPZ, 2929.65±127.92 vs. Control, 2864.29±123.44; p=0.5477); 5+4w (3400.91±232.62 vs. Control, 3012.23±256.68; p=0.1). For *Mog-*positive cell counts (in cells/mm^2^): 1w (CPZ, 28.86±9.66 vs. Control, 1485.37±160.07; p<0.0001); 2w (CPZ, 19.77±10.24 vs. Control, 1245.21±148.46; p<0.0001); 3w (CPZ, 336.98±132.64 vs. Control, 1376.08±157.04; p=0.0002); 4w (269.55±37.20 vs. Control, 1231.21±137.22; p<0.0001); 5w (250.24±41.15 vs. Control, 1439.50±183.75; p=0.0009); 5+1w (CPZ, 1674.099±186.88 vs. Control, 1296.28±77.09; p=0.0177); 5+2w (CPZ, 1775.62±107.35 vs. Control,1360.74±76.45; p=0.0112); 5+3w (CPZ, 1700.13±150.17 vs. Control, 1355.93±56.43; p=0.0099); 5+4w (CPZ, 1829.02±129.01 vs. Control, 1370.25±128.05; p=0.0047).

**Fig. 4.**
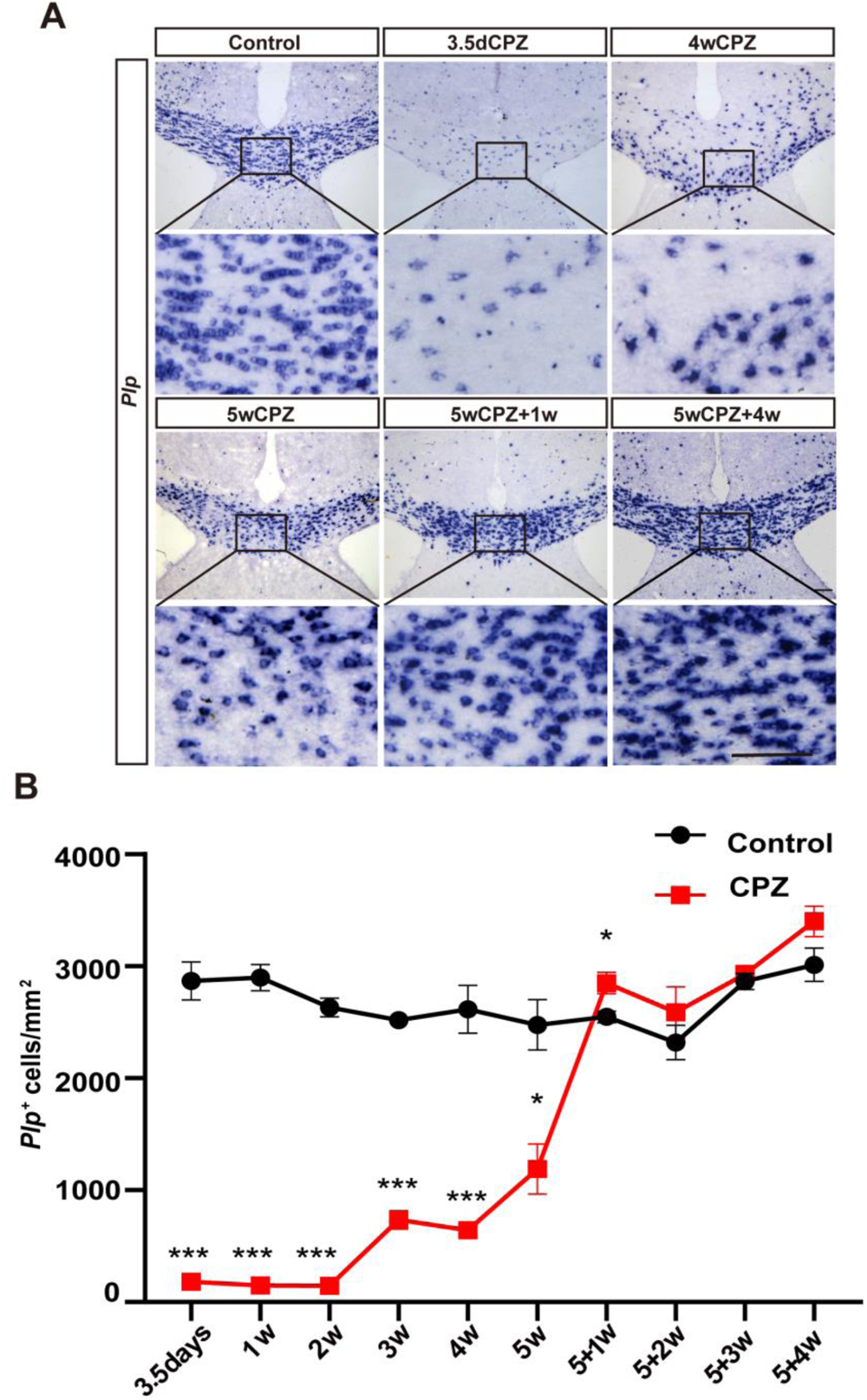
Dynamics of *Plp*-positive mature OLs in standard CPZ model. (A) *In situ* hybridization with *Plp* specific probe. Representative images of control, CPZ-treated mice at 3.5d, 4w, 5w; and 1, 4 weeks after CPZ withdrawal. (B) Graph showing quantification of *Plp+* cells at different time points. Control group (black line); treatment group (red line). (3.5d, n=5/group; 1, 2w, 4w, 5+1w, 5+3w, 5+4w, n=4/group; 3w, 5w, 5+2w, n=3/group). Data are mean± SEM. Asterisks indicate significant difference from normal chow mice. \**p*<0.05; \*\**p*<0.01; \*\*\**p*<0.001. Two-tailed unpaired t test. Scale bars: 100um. CPZ, cuprizone; *Plp*, proteolipid protein gene, marker for mature oligodendrocytes.

**Fig. 5.**
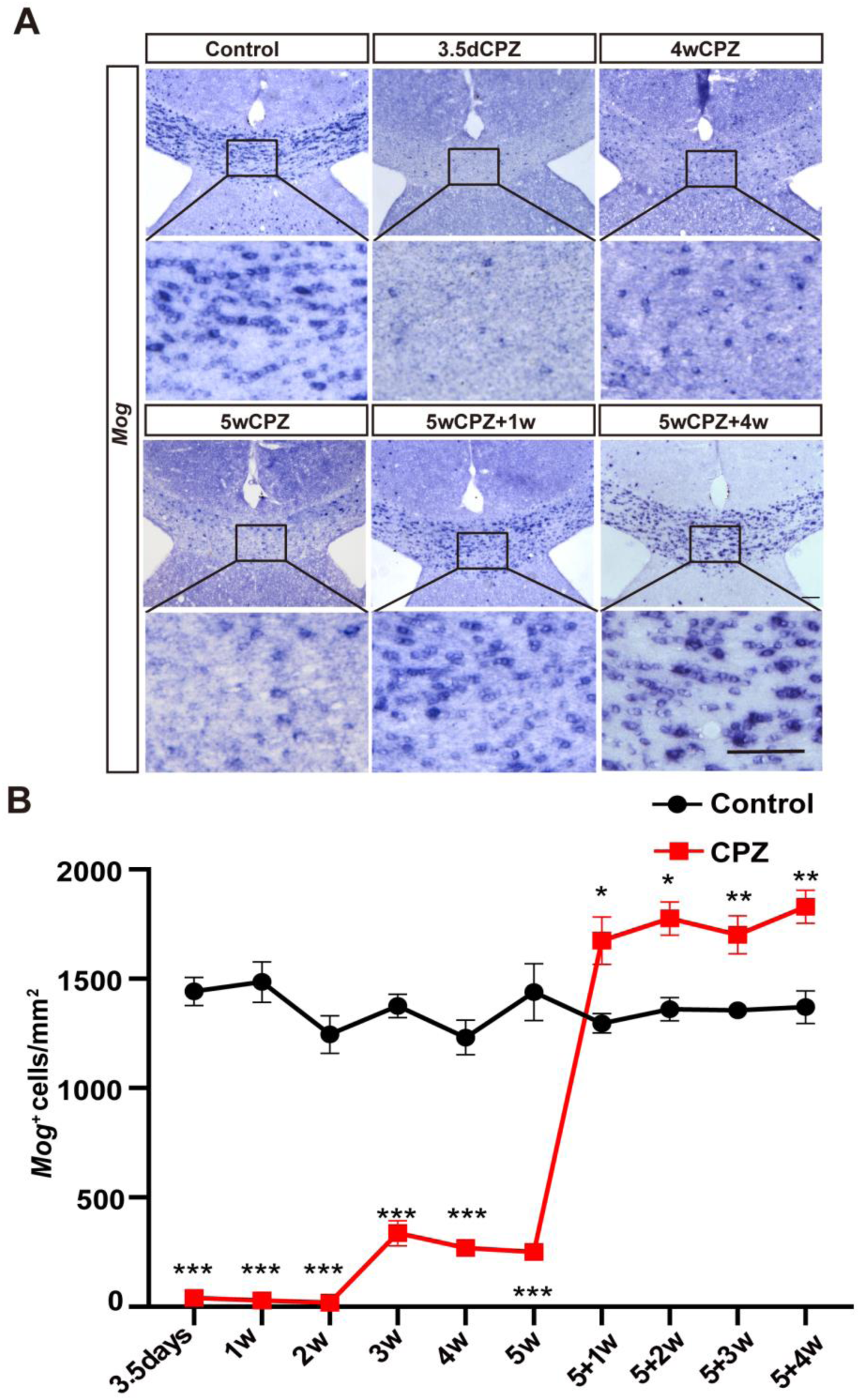
Dynamics of *Mog*-positive mature OLs in standard CPZ model. (A) In situ hybridization with *Mog* specific probe. Representative images from control, CPZ-treated mice 3.5d, 4w, 5w; and 1, 4 weeks after CPZ withdrawal. (B) The graph shows the quantification of *Mog^+^* cells at different time points. Control group (black line); the treatment group (red line). (3.5d, n=5/group; 1, 2w, 4w, 5+1w, 5+3w, 5+4w, n=4/group; 3w, 5w, 5+2w, n=3/group). Data are mean± SEM. \**p*<0.05; \*\**p*<0.01; \*\*\**p*<0.001. Asterisks indicate significant difference from mice on the normal chow. Two-tailed unpaired t test. Scale bars: 100um. CPZ, cuprizone; *Mog*, myelin oligodendrocyte glycoprotein gene, marker for mature oligodendrocytes.

In the case of *Mbp* ISH analysis, consistent with our previous report (Xiao et al., 2016), two types of *Mbp-*positive cells were recognized. Type 1 were *weak-Mbp^+^* OL cells with fade hybridization signal located only in the cytoplasm of the cell body, representing normal old mature OLs similar to these signals of *Plp* and *Mog*, which accounted for the majority (>95%) of all *Mbp^+^*OL cells. Type 2 were strong *Mbp^+^* OL cells with significantly higher hybridization signal density in the cell body than type 1 cells, and the signal also extended to the cell processes, giving a spider-like morphology. These two types of *Mbp^+^*OLs are clearly distinguishable in the CC and even more obvious in the cortex (Fig.6A, Fig.S2). Analyzing both types of *Mbp-*positive cells as a whole in terms of signal density, we can see a very similar dynamic time curve to that of *Plp-* and *Mog-*positive cell numbers (Fig.6B). When counting the number of type1 *weak-Mbp^+^* OL cells, there showed a similar cliff-like decline as the number of *Plp-* and *Mog-*positive cells from as early as 3 days after CPZ challenge. However, the rate of return to control level is much slower. The gradual increase during the late demyelinating phase (3-5 weeks) and the sharp increase in early remyelination phase (5+1) that occurs *Plp-* and *Mog-*positive cells were not observed. There was a gradual increase only after 5 weeks, when CPZ was withdrawn, and it still remained below the control level for at least 4 more weeks (Fig.6D). For detailed cell counts (in cells/mm^2^): 3.5d (CPZ, 76.84±7.32 vs. Control, 1479.4±269.82; p<0.0001); 1w (CPZ, 54.05±14.26 vs. Control, 1519.49±238.09 ; p<0.0001); 2w(CPZ, 32.94±6.99 vs. Control, 1485.97±124.49; p<0.0001),3w(CPZ, 124.63±74.49 vs. Control, 1833.74±212.05; p<0.0001); 4w (CPZ, 143.17±45.98 vs. Control, 1758.17±289.66; p<0.0001); 5w (CPZ, 178.38±17.19 vs. Control, 1570.64±120.54; p<0.0001); 5+1w (CPZ, 385.31±34.75 vs. Control, 1702.86±277.47; p=0.0002); 5+2w (CPZ, 997.95±77.90 vs. Control, 1649.46±281.41; p=0.0343); 5+3w (CPZ, 1145.62± 201.11 vs. Control, 1518.42±159.81; p=0.0457); 5+4w (CPZ, 1491.45±195.47 vs. Control, 1638.31±101.84; p=0.2923)).

**Fig. 6.**
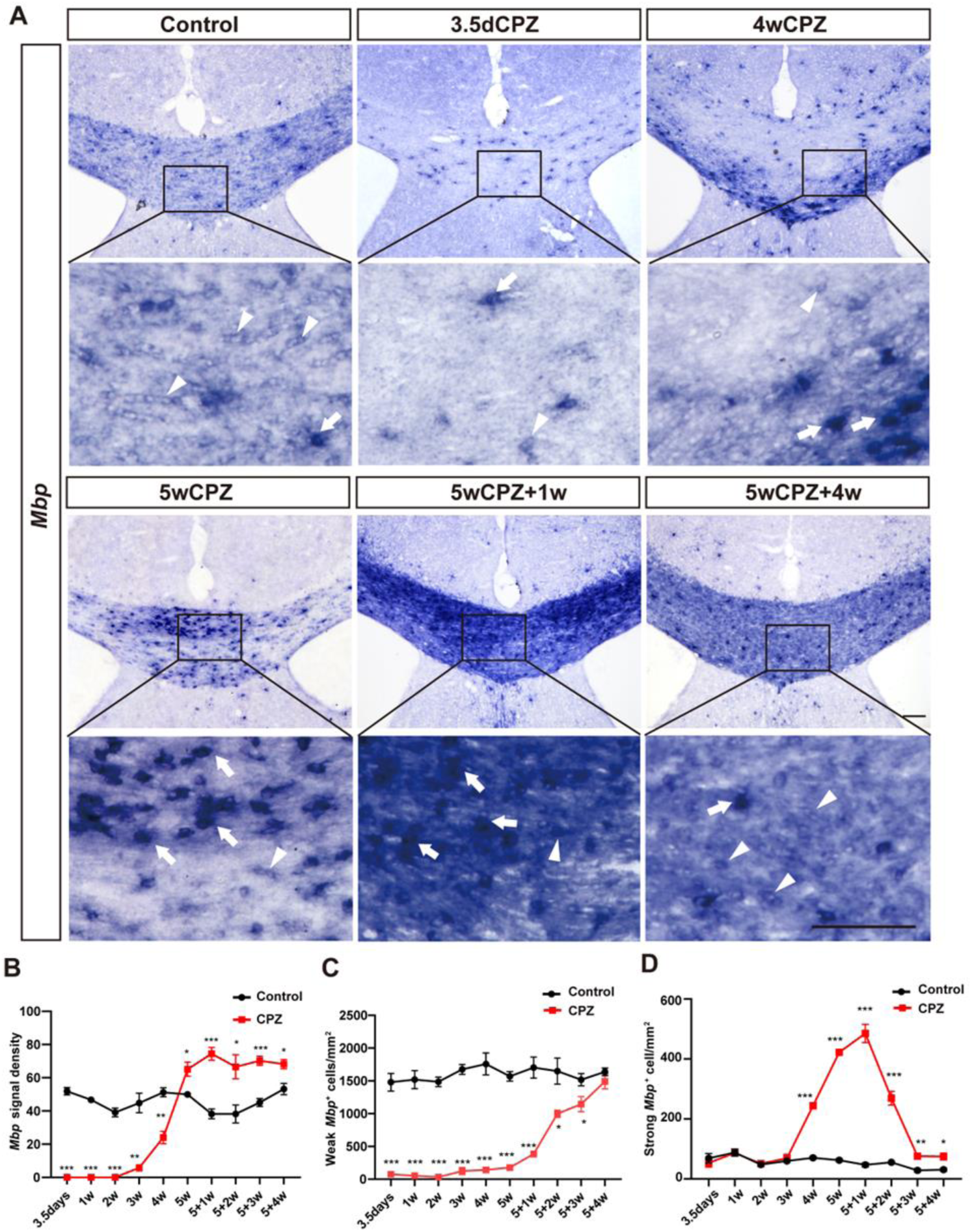
Dynamics of *Mbp*-positive cells in standard CPZ model. (A) *In situ* hybridization with *Mbp* specific probe. Representative images of control, CPZ-treated mice at 3.5d, 4w, 5w; and 1, 4 weeks after CPZ withdrawal. Arrows indicate the strong *Mbp*^+^ positive cells, whereas arrow heads indicate the weak *Mbp^+^* positive cells. (B) Graphs show the quantification of total *Mbp^+^* signal density at different time points. (C) Graphs show the quantification of weak *Mbp^+^*cells at different time points. (D) Graphs indicate the quantification of strong *Mbp^+^* cells at different time points. Control group (black line); the treatment group (red line). (3.5d, n=5/group; 1, 2w, 4w, 5+1w, 5+3w, 5+4w, n=4/group; 3w, 5w, 5+2w, n=3/group). Data are mean± SEM. \**p*<0.05; \*\**p*<0.01; \*\*\**p*<0.001. Asterisks indicate significant difference from mice on the normal chow. Two-tailed unpaired t test. Scale bars: 100um. CPZ, cuprizone; *Mbp*, myelin basic protein gene, marker for mature oligodendrocytes.

Thus, in consistent with the CC1 immuno-staining, these ISH results also clearly demonstrated a quite rapid response, i.e. loss of mature OLs to the CPZ challenge, followed by a reversed and sustained increase of these cells during the remyelination period lasting at least for 4 weeks after CPZ withdrawal.

### Dynamics of newly-forming/formed OLs

The differentiation of OPCs to generate new OLs and produce new myelin are essential processes for successful remyelination in human demyelinating diseases such as MS and in animal models of demyelination, although some recent studies have also argued for the contribution of pre-existing surviving mature Ols (Bacmeister et al., 2020; Crawford et al., 2016; Duncan et al., 2017; Neely et al., 2022). Therefore, it is important to determine the profile of new OL genesis throughout the course of the current standard CPZ model. As previous suggested, the above-mentioned spidery strong-*Mbp^+^*(type 2) cells are actually the newly-forming/formed OLs that are largely overlapped with the population of *Enpp6* positive cells (Xiao et al., 2016). These cells were then counted. It showed a strong increase from the 4^th^ week of CPZ feeding (4w (CPZ, 243.80±17.53 cells/mm^2^ vs. Control,140.79±8.37 cells/mm^2^; p<0.0001)), which peaked (∼10 fold of control) at 1 week after CPZ withdrawal (CPZ, 485±52.8 cells/mm^2^ vs. Control, 46.70±11.87 cells/mm^2^; p<0.0001), and declined sharply over the following 2 weeks, but still remained significantly higher than control (∼2 fold of control) (Fig. 6B). For detailed cell counts in other time points (in cells/mm^2^): 3.5 days (CPZ, 53.36±28.61 vs. Control, 68.57±31.63; p=0.4960);1w (CPZ, 86.39±3.47 vs. Control, 87.77±20.92; p=0.9138); 2w (CPZ, 50.66±15.57 vs. Control, 47.96±9.47; p=0.8507); 3w (CPZ, 68.69±17.86 vs. Control, 59.71±4.36; p=0.5276); 5w (CPZ, 422.50±4.21 vs. Control, 62.30±7.41; p<0.0001); 5+2w (CPZ, 269.04±32.17 vs. Control, 55.21±3.46; p=0.0007); 5+3w (CPZ, 75.6±11.82 vs. Control, 28.44±9.72; p=0.0018); 5+4w (CPZ,74.36±20.04 vs. Control, 30.91±9.64; p=0.0148).

To further characterize the feature of new OL genesis during whole course of this model, we further took the advantage of our recently identified OL lineage cell stage-specific marker, ENPP6, a choline-specific ecto-nucleotide pyrophosphatase /phosphodiesterase (ENPP), which is highly expressed in newly-forming/formed OLs and at a much lower levels in these more mature/old (earlier-formed) myelinating OLs (to the extent that the ISH signal would easily disappear if the development time were shortened), but not at all in OPCs, neurons, astrocytes or vascular endothelial cells (Xiao et al., 2016; Zhang et al., 2014). As shown in figure 7A, the result showed a quite similar dynamic curve of *Enpp6^+^* cells as the type 2 spidery strong-*Mbp^+^* cells, while two main differences are also evident. Firstly, there was a rapid and significant an increase (∼51%) in *Enpp6^+^* cells from 3.5 days after CPZ intake (CPZ, 141.63±8.14 cells/mm^2^ vs. Control, 93.81±11.31 cells/mm^2^; p<0.0001), suggesting an early response as seen in CC1^+^, *Pdgfra*^+^, *Plp^+^*, *Mog*^+^ and weak-*Mbp*^+^ cells. Secondly, the number of *Enpp6^+^* cells peaked at 5 weeks after CPZ (CPZ, 353.1±55.21 cells/mm^2^ vs. Control, 48.95±6.48 cells/mm^2^; p=0.0015), which was one week earlier than that of spidery strong-*Mbp^+^*cells, while also remaining at a higher level than control (∼2 fold of control) at least until 4 weeks after CPZ withdrawal (CPZ,101.52±24.54 cells/mm^2^ vs. Control, 45.63±8.99 cells/mm^2^; p=0.01) (Fig. 7B). For detailed cell counts in other time points (in cells/mm^2^): 1w (CPZ,126.65±12.65 vs. Control, 90.93±16.44; p=0.0246); 2w (CPZ, 123.52±26.8 vs. Control, 68.06±11.9; p=0.0265),3w (CPZ, 130.65±6.13 vs. Control, 63.87±1.23; p=0.0001); 4w (CPZ, 203.52±21.55 vs. Control, 57.36±13.03; p<0.0001); 5+1w (CPZ, 282.17±41.29 vs. Control, 60.77±8.87; p=0.0001); 5+2w (CPZ, 158.43±10.16 vs. Control, 41.95±3.78; p=0.0001); 5+3w (CPZ,125.01±25.78 vs. Control, 51.21±6.67; p=0.003)

**Fig. 7.**
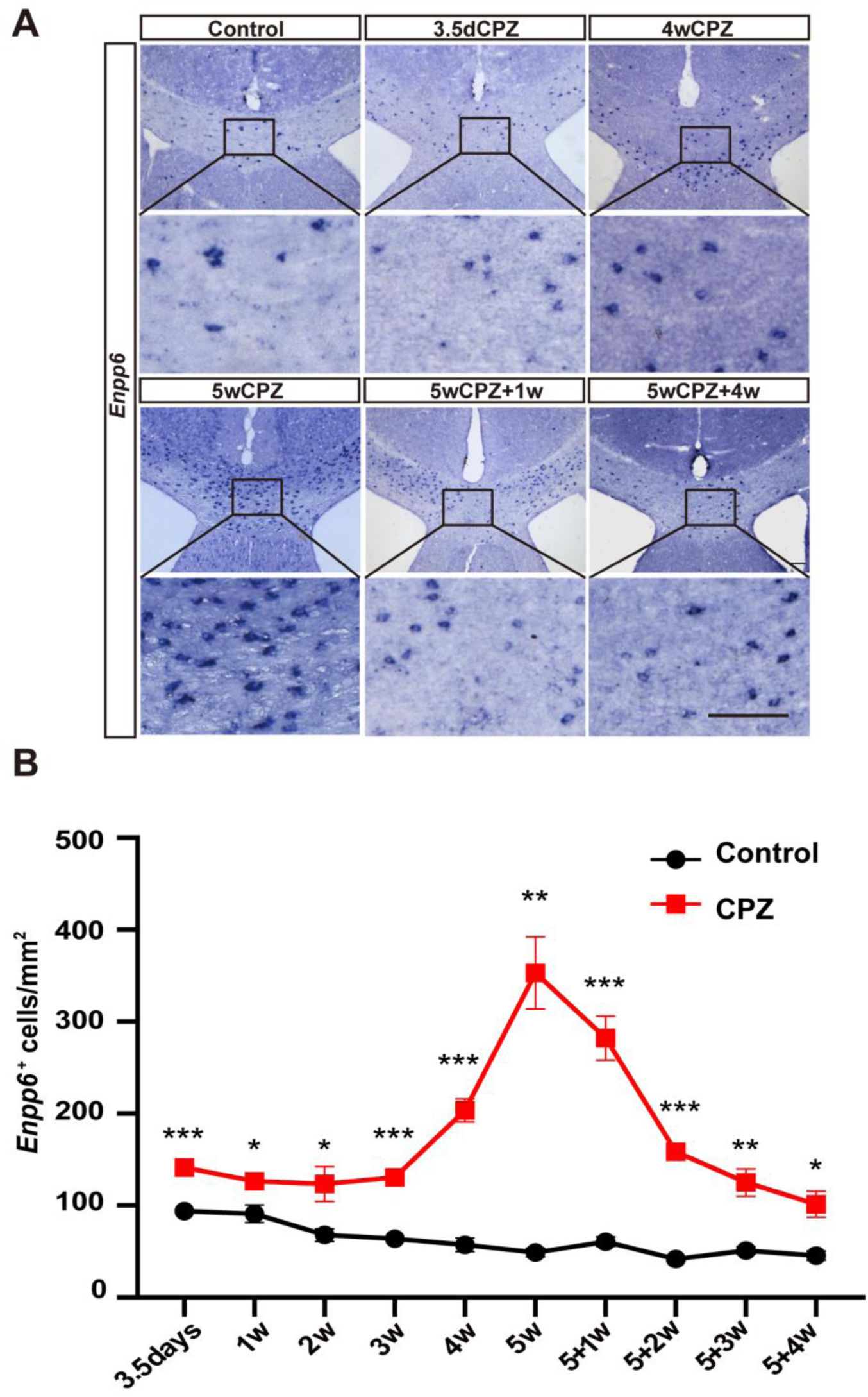
Dynamics of *Enpp6*-positive newly-forming/formed OLs in standard CPZ model. (A) *In situ* hybridization with an *Enpp6* specific probe revealed the number of newly forming/formed oligodendrocytes in the corpus callosum. Representative images of control, CPZ-treated mice 3.5d, 4w, 5w; and 1, 4 weeks after withdrawal of CPZ. (B) Graph shows the quantification of *Enpp6*^+^ cells at different time points. Control group (black line); the treatment group (red line). (3.5d, n=5/group; 1, 2w, 4w, 5+1w, 5+3w, 5+4w, n=4/group; 3w, 5w, 5+2w, n=3/group). Data are mean± SEM. \**p*<0.05; \*\**p*<0.01; \*\*\**p*<0.001. Two-tailed unpaired *t* test. Scale bars: 100um. CPZ, cuprizone; *Enpp6*, ectonucleotide phosphodiesterase/pyrophosphatase 6 gene, newly forming/formed oligodendrocyte marker

## Discussion

Since the first report of the neurotoxicity of the copper chelator cuprizone in Swiss albino mice by Carlton in 1966 (Carlton, 1966) and the subsequent early description of CPZ-induced OL death and demyelination in the superior cerebellar peduncles of ICI mice (Blakemore, 1972), the CPZ-induced demyelination model has been intensively studied and optimized for more than five decades (Vega-Riquer et al., 2019). Studies in different species (e.g. mice, rats, guinea pigs, hamsters) and strains (e.g. Swiss, C57BL/6, BALB/cJ, ICI, CD1, SJL mice) at different dietary doses ranging from 0.1% to 5% revealed a highly species- and strain-dependent susceptibility to CPZ-induced demyelination, and the extent of demyelination also varied with dose, age, sex and brain region (Skripuletz et al., 2011; Vega-Riquer et al., 2019; Zirngibl et al., 2022). It is therefore difficult to compare data between different mouse strains, and each model needs to be characterized before firm conclusions can be drawn.

The now days mostly used standardized CPZ mice model was first established in 1998 by the Matsushima group (Hiremath et al., 1998). In that study, the authors used young adult (8-10 weeks old) male C57BL/6 mice and performed a dosage screen (from 0.1% to 0.6%) by examining the demyelination in the corpus callosum of cerebral white matter, a brain region where the extent of demyelination can be scored more easily and consistently than the previously described superior cerebellar peduncles with LFB-PAS staining (Skripuletz et al., 2011). They found that at the dose of 0.2% for 6 weeks, CPZ could induce complete demyelination in the corpus callosum while having no detectable toxic effect on the peripheral organs such as the liver and causing only mild weight loss in the mice. Subsequent studies have also shown significant of demyelination in the grey matter of the cerebrum and cerebellum (Gudi et al., 2009; Koutsoudaki et al., 2009; Pott et al., 2009), although the corpus callosum remains to be the most commonly analyzed region. Moreover, a 5-week feeding regimen was found to be more suitable than the initial 6-week regimen for the acute demyelination model, since demyelination in the corpus callosum had already peaked/completed at this time point, and detectable remyelination was observed at 6 weeks even under the continuous feeding of CPZ (Matsushima and Morell, 2001; Zirngibl et al., 2022). The introduction and standardization of the CPZ model in C57BL/6 mice has made it possible to share and compare data from different laboratories to gain a more reliable insight into the molecular and cellular mechanisms underlying de- and remyelination, with particular facilitation for the use of various transgenic mice, which in most circumstances bear a C57BL/6 genetic background. Thus, the more detailed information we can get about this standardize model, the better it can be used as a model for human demyelinating diseases.

In this study, we have carried out a whole-process mapping of the OL lineages cell dynamic in the corpus callosum region during the de- and remyelination period in the standardized CPZ model with the dose of 0.2% for 5 weeks and followed by up to 4 weeks’ recovery in the young adult male C57BL/6 mice. In particular, we used a series of ISH probes targeting the well-known marker genes for different stages of OL lineage cells such as *Pdgfra, Plp, Mbp, Mog,* as well as a recently reported maker for newly forming/formed OLs, namely *Enpp6*, to access a reliable counting of these cells. To our knowledge, this type of survey has never been conducted before.

Our results showed a rather unexpectedly rapid response of the OL lineage cells to the CPZ challenge. ISH analysis revealed an almost complete disappearance of *Plp-* and *Mog-*positive (to the level of ∼6% and ∼3% of control respectively) and weak *Mbp-*positive (-5% for control) mature OLs only 3.5 days after the onset of CPZ feeding. Although the Matsushima group had already observed a profoundly depressed expression levels of mRNA for several myelin related genes, including *Mbp,* in the CC region after 1 week of CPZ exposure by Northern blotting analysis of isolated brain tissue when they initially established this standardized model (Morell et al., 1998), the current ISH results on brain sections were still quite a surprising finding for us. It is difficult to judge for sure whether the early drastic decrease in *Plp-* and *Mog-*positive cells implies a drastic death/loss of the mature OLs or just an extreme-down regulation of the expression of these myelin-associated genes by the OLs while they themselves are still alive. However, as our immunofluorescence staining also showed a rapid (at 3.5 days) and dramatic (∼40%) decrease in the number of CC1-positive cells, another protein marker of mature OLs, an early onset of OL death is more likely. These results are in line with at least two previous studies. In the first study, the authors reported the appearance of dying OLs expressing activated caspase 3 which was accompanied by a significant decrease in NogoA-positive mature OLs after 6 days of CPZ feeding at the dose of 0.3%, and they also claimed that apoptotic OLs were present not only in the corpus callosum but also in the cortex 2 days after the start of the CPZ diet, although they did not show the data (Hesse et al., 2010).The second and more recent study showed the loss of CC1-positive OLs in the corpus callosum as early as 2 days after CPZ exposure in the standardized CPZ model (Jhelum et al., 2020). Therefore, our data, together with others, strongly suggest that CPZ can trigger OL death well before the time when demyelination is detectable, which is 2-3 weeks after CPZ administration in the standard model, though the detail mechanisms for CPZ induced OL death are still unknown (Zirngibl et al., 2022).

Interestingly, the early (3.5 days) slight (∼0.5-fold more than controls, while the peak was ∼7-fold) but significant increase in *Enpp6*-positive newly-forming/formed OLs was accompanied by a concomitant and very comparable degree of decrease (∼52%) in *Pdgfra*-positive OPCs, with the specific values of ∼50 per mm^2^ increase for *Ennp6*-positive cells, and ∼80 per mm^2^ decrease for *Pdgfra*-positive cells. It is very likely that most of these 80 *Pdgfra*-positive OPCs have progressed to *Enpp6*-positive newly forming/formed OLs, with the reaming ∼30 cells to the *Enpp6* and *Pdfgra* double-negative stage (Xiao et al., 2016). The continuous increase in both *Pdfgra*- and *Enpp6*-positive cells implicated a very rapid response/mobilization of OPC to OL death and that OPCs and newly-forming/formed OLs are both resistant to CPZ toxicity in vivo, which is in direct contrast to the mature OLs. In vitro, CPZ can lower the mitochondrial transmembrane potential of cultured OLs without affecting microglia, astrocytes, and neurons (Bénardais et al., 2013), but it does not it kill the Ols (Pasquini et al., 2007). Copper is a trace element and a critical component of many mitochondrial metalloenzymes in which it acts as an electron donor or acceptor (Stern et al., 2007), and these enzymes, such as monoamine oxidase (Kesterson and Carlton, 1971) and cytochrome c (Cyt c) oxidase (Acs et al., 2013; Venturini, 1973), are critical for oxidative phosphorylation and energy production. Thus, it is generally accepted that the neurotoxicity of CPZ is mainly induced by a disturbed mitochondrial function and energy metabolism (Kipp et al., 2009), although whether these consequences are a direct result of copper chelation remains unclear, as copper supplementation has a limited ability to reduce CPZ toxicity (Carlton, 1967). However, it is still a mystery why CPZ is specifically toxic to mature OLs without damaging other types of neural cells in the brain (Bénardais et al., 2013; Zirngibl et al., 2022) We suspect that the high energy requirement of OLs to maintain their myelin sheaths may be one reason why they extremely susceptible to CPZ. The survival and activation/proliferation of OPCs, as well as and their enhanced and continuous differentiation into *Enpp6*-positive newly-forming/formed OLs, despite the late onset (from 2-3 weeks and beyond) demyelination under the continuous pressure of CPZ, underpinned a strong potential for remyelination after CPZ withdrawal. The sequential peaking of *Pdfgra*-positive cells at 4 weeks, *Enpp6*-positive cells at 5 weeks, and strong *Mbp*-positive cells at 5+1 weeks, and these cells together with the *Plp*-, weak Mbp-, and *Mog*-positive cells that have decreased sharply at early time and gradually recovered from 3 weeks to 5 or 5+1 weeks, respectively, and then remained at a higher level than intact control as long as to 5+4 weeks, provided a clear overall dynamic profile of OL lineage cells throughout de-and remyelination period of the standardized CPZ model (Fig.8). The prolonged presence of these cells on at a relatively higher level suggested a still ongoing remyelination process that requires the continuous generation of new OLs and the synthesis of myelin-related proteins for the forming of new compact myelin, although the myelin score by LFB-PAS staining has already returned to a level indistinguishable from the normal control by 3 weeks after CPZ withdrawal.

**Fig. 8.**
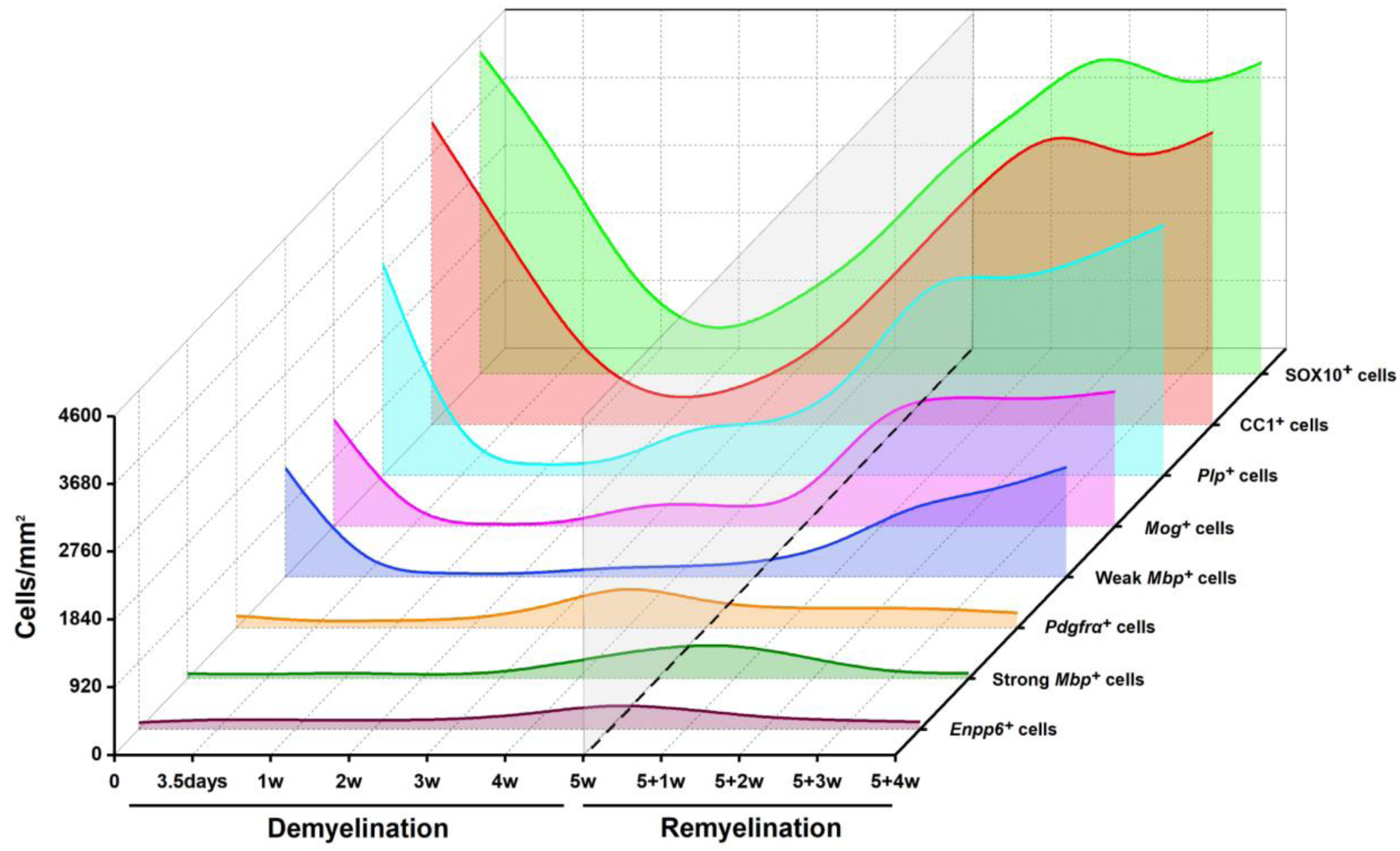
Rapid and prolonged response of oligodendrocyte lineage cells in standard acute CPZ demyelination model. Relative cell number dynamics of the OL lineage by different markers during the de- and remyelination phases of the standard CPZ model were shown.

In summary, the CPZ model has been shown to mimic many aspects of human demyelinating diseases such as MS, and to be a useful experimental approach to study these diseases and test potential therapies, especially after its standardization on C57BL/6 mice (Vega-Riquer et al., 2019; Zirngibl et al., 2022). In the present study, we demonstrated an early death of mature OLs upon CPZ challenge and a rapid response of OPCs to this death, as well as a sustained increase in the number of myelin-producing new OLs in the standardized CPZ model. These data shed new light on the detailed cell dynamics underlying this model and may serve as a basic reference system for future studies of the effects of any intervention on de- and remyelination using the CPZ model. It may also indicate the need to optimize the timing window for pro-remyelination therapies in demyelinating diseases such as MS according to the intrinsic cell dynamics.

## Data availability

Data will be made available on request.

## Acknowledgments

We express our thanks to professor Huiliang Li (Wolfson Institute for Biomedical Research, UCL) for his helpful discussion of the project, critical reading and kind suggestions to the manuscript. This work was supported by STI 2030-Major Project 2021ZD0201703; National Natural Science Foundation of China (31970913 and 32170957); Guangdong Basic and Applied Basic Research Foundation (2021A1515012156); and the Key-Area Research and Development Program of Guangdong Province (2019B030335001)

## Supplemental Figure

**Supplementary Fig. 1.**
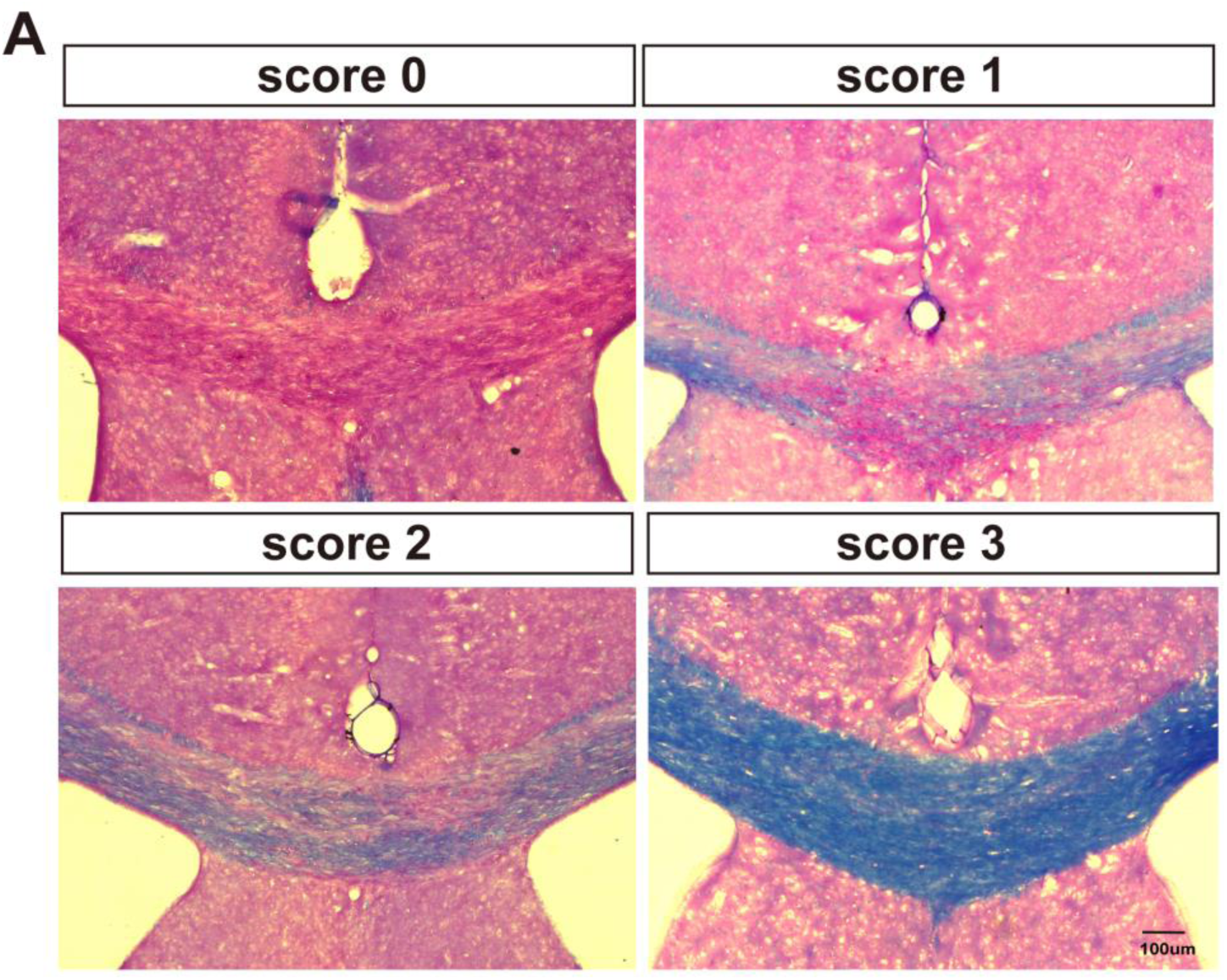
Representative plots for each score of myelin by LFB staining. Scale bar: 100 um

**Supplementary Fig. 2.**
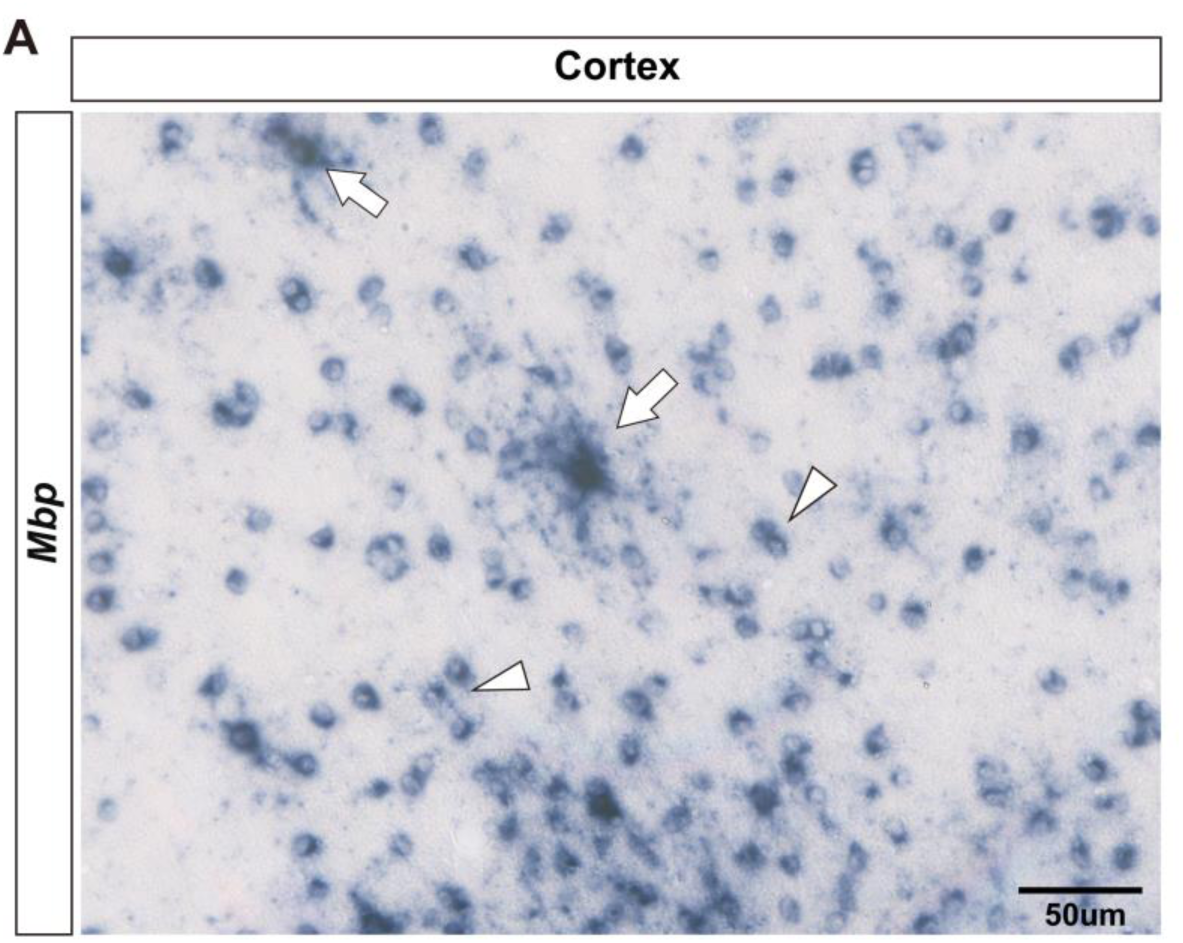
Representative images for strong (arrows) and weak (arrow heads) *Mbp^+^* cell in cortex. Scale bar: 50 um

